# In-field capable loop-mediated isothermal amplification detection of fall army worm (*Spodoptera frugiperda*; Lepidoptera: Noctuidae) larvae using a rapid and simple crude extraction technique

**DOI:** 10.1101/2021.08.09.455740

**Authors:** B. S. Congdon, C.G. Webster, D. Severtson, H. Spafford

## Abstract

Fall armyworm, *Spodoptera frugiperda* (Lepidoptera: Noctuidae), is an economically important pest worldwide and has recently been identified in Australia. Morphological identification of *S. frugiperda* at early larval stages can be difficult often requiring expert microscopy analysis. Rapid and accurate in-field diagnosis is vital for management decision support and there are no tools currently available for this purpose. In this study, a sensitive, specific and in-field capable loop-mediated isothermal amplification (LAMP) assay was developed to detect *S. frugiperda* larvae. A primer set based on a highly conserved region of the *S. frugiperda* cytochrome oxidase subunit 1 (COX1) gene provided detection within 30 min from both total DNA and crude extractions. The crude extraction technique of crushing 10 mg of *S. frugiperda* material in 50 μL ddH_2_0 and further diluting the homogenate in ddH_2_0 is rapid, simple and does not require heat blocks, centrifuges or special buffers increasing its utility as a field-based technique. The primer set detected as little as 24 pg of *S. frugiperda* DNA and did not cross-react with any other of the lepidopteran species tested that are easily confused with *S. frugiperda* in Australia. Therefore, this assay could be used in-field to correctly identify the presence of *S. frugiperda* and thereby greatly assist with timely management decisions.

## Introduction

Fall armyworm, *Spodoptera frugiperda* J.E. Smith (*Lepidoptera: Noctuidae*), is an economically important pest worldwide that can cause significant yield losses in grains and horticultural crops, pastures, rangelands and urban landscapes (Cruz and Turpin 1983, Buntin 1986, Pantoja et al. 1986, Chamberlin and All 1991, Chimweta et al. 2020, Overton et al. 2021). *S. frugiperda* was first detected in Australia in early 2020 on two Torres Strait islands before spreading to northern regions of Queensland, the Northern Territory, and Western Australia (Queensland Government 2020) in which it rapidly established in crops such as maize/sweet corn (*Zea mays* L.) and sorghum (*Sorghum bicolor* L. Moench), and Rhodes grass (*Chloris gayana* Kunth). As predicted, *S. frugiperda* has migrated further south, as far as Tasmania, into key agricultural production areas in which it poses a risk to summer crops such as sweetcorn, maize and sorghum and winter cereal production (Maino et al. 2021). In addition to its preference for grasses, *S. frugiperda* is noted to feed on over 350 plant species including other economically important crops such as cotton (*Gossypium hirsutum*), peanut (*Arachis hypogaea*), chickpea (*Cicer arietinum*), soybean (*Glycine max*), vegetables and some fruit crops also grown in northern Australia and other parts of the country (Montezano et al. 2018).

In its broad host range among Australian crops, there are a number of congeneric and other lepidopteran species which occur sympatrically with *S. frugiperda*. Many of the noctuid moths in particular are very similar in appearance to *S. frugiperda*, especially in early larval instars. The shared characteristics between species make identification very difficult because either visual identification by trained entomologists or diagnosis by laboratory-based PCR tests is required (e.g. Jing et al., 2020). Rapid and accurate identification of *S. frugiperda* is most necessary to detect the presence of a larval population in a crop before it causes significant damage. Accurate identification is even more crucial when one or more species present in the crop carry insecticide resistance alleles to ensure insecticide efficacy and assess the appropriateness of other control measures such as use of nuclear polyhedrosis virus. Therefore, accurate and rapid in-field diagnostic tools are essential to provide timely decision support to growers without having to send samples to a laboratory.

Loop-mediated isothermal amplification (LAMP) has been used in tandem with simple and inexpensive crude extraction techniques easily adapted to the field for detection of insects, pesticide resistance mutations, plant pathogens and more (e.g. Congdon et al., 2019; Sial et al. 2020; Pusz-Bochenska et al. 2020; Przybylska et al. 2015). Such an approach offers a promising avenue for regional based and in-field testing of larvae infesting local crops for presence of *S. frugiperda*. This study describes the development of a LAMP assay with a simple crude extraction technique for detection of *S. frugiperda*.

## Materials and Methods

### Specimen collection

*S. frugiperda* larvae used to provide positive controls for the LAMP assay were taken from a colony maintained at the Frank Wise Institute for Tropical Agriculture (Kununurra, Western Australia). All other larvae specimens were collected from field sites around Western Australia.

### Total DNA and crude extraction

Total DNA was extracted from insect tissue using a DNeasy®Blood & Tissue Kit according to manufacturer instructions (QIAGEN, Australia). Crude extractions were conducted using a polypropylene pellet pestle (Sigma-Aldrich, USA) to grind 10 mg of larval abdominal tissue in a 1.5 mL microcentrifuge tube containing 50 μL ddH_2_0 for five to ten seconds. Then, 10 μL of the resulting homogenate was diluted in 1 mL of ddH_2_0. Crude extracts of larvae were either used as template immediately or frozen at −20°C until required.

### LAMP assay development

#### Primers

An alignment of eight *S. frugiperda* mitochondrial genomes obtained from GenBank (Table 1) was made using Geneious Prime 2019.2.3 (Biomatters, New Zealand) and the consensus sequence was used to generate several LAMP primer sets using PrimerExplorer V5 software (available at http://primerexplorer.jp/lampv5e/index.html) with default settings. The primer set FAW-1.2, derived from the cytochrome c oxidase subunit I (COX1) gene nt sequences, was selected for further testing (Table 2).

**Table 1.** Eight *Spodoptera frugiperda* mitochondrial genomes obtained from GenBank used for loop-mediated isothermal amplification primer development

| Accession No. | Country | Author |
| --- | --- | --- |
| MN803322 | Benin | Kim et al. unpublished |
| KU877172 | Brazil | Paula et al. unpublished |
| KM362176 | China | Liu et al. 2016 |
| MN599980 | Nigeria | Kim et al. unpublished |
| MN385596 | South Korea | Seo et al. 2019 |
| MN599983 | South Korea | Kim et al. unpublished |
| MN599981 | South Korea | Kim et al. unpublished |
| MN599982 | South Korea | Kim et al. unpublished |

**Table 2.**
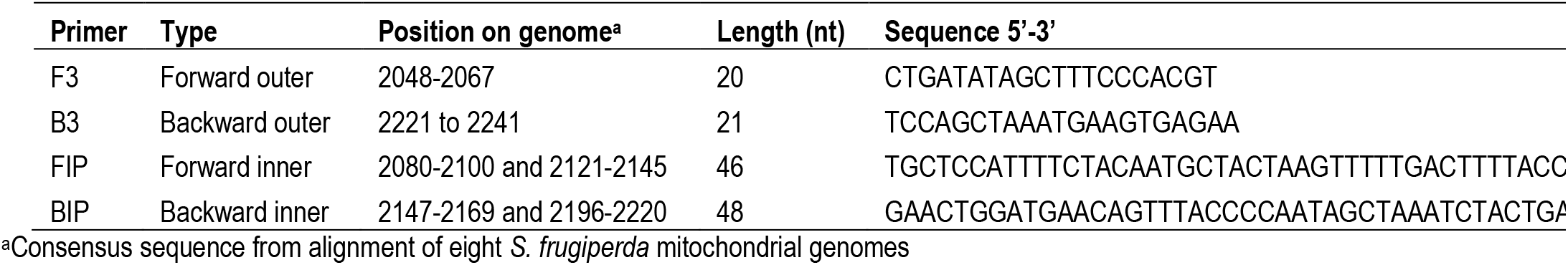
*Spodoptera frugiperda*-specific loop-mediated isothermal amplification primer set FAW-1.2

#### Assay

All LAMP reactions were done using a dual-block Genie® II instrument or single-block Genie®III (Optigene, United Kingdom). In a total volume of 25 μL, the reaction mixture always contained 15 μL ISO-004 master mix (Optigene, United Kingdom), 1 μL total DNA extraction or 3 μL diluted crude extraction template, primer mix consisting of a 1:4 ratio of F3 and B3:FIP and BIP and ddH_2_0. Each set of eight reactions generally included a water (negative control) and at least one total DNA positive control.

#### Parameter selection

To select the parameters for the LAMP assay, primer set FAW-1.2 was used to test total DNA and crude extractions of *S. frugiperda, S. litura* and *H. punctigera* in duplicates at three amplification temperatures (63°C, 65°C and 67°C), and two primer concentrations; 12.5 pmol (5.6 μL primer mix per reaction) or 18.5 pmol (8.2 μL). The assay was run for 30 min followed by a 10 min annealing step. The time to 10,000 fluorescence, peak fluorescence and annealing temperature were recorded. This process was repeated twice more.

#### Diagnostic sensitivity and specificity

The type I (false positive) and type II (false negative) error rate of the assay was assessed by testing two crude extractions each of *S. frugiperda, H. punctigera* and *Plutella xylostella* in duplicates. This process was repeated eighteen times for a total of 72 known positives and 72 known negatives.

For future testing, a sample was considered positive if fluorescence exceeded 10,000 within 20 min, reached a peak flourescence of over 20,000 and had an annealing temperature within 1°C of the positive control. These standards are specific to this assay.

#### Analytical sensitivity

Three total DNA extractions of *S. frugipera* larvae were used to assess sensitivity of the LAMP assay. These extractions had 37, 240 and 290 ng/μL of DNA, respectively, as assessed using a NanoDrop^TM^ 2000 spectrophotometer (Thermo Fisher Scientific, USA). Each extraction was tested by LAMP in a series of ten-fold dilutions up to a 10^−6^ dilution and 1 μL of each dilution was used as template. The time to 10,000 fluorescence, peak fluorescence and annealing temperature were recorded. The assay was run for a further 30 minutes (total of 1 h) to assess any late amplification of low concentration reactions. This process was repeated twice more.

#### Analytical specificity

Primer specificity was determined by testing primer set FAW-1.2 against total DNA and crude extractions of *Spodotera frugiperda, Spodoptera mauritia, S. litura, Chrysodeixis eriosoma, C. subsidens, Dasygaster padockina, Helicoverpa armigera, H. punctigera, Heliothis punctifera, Hellula hydralis, Hyles livornicoides, Leucania loreyi, L. stenographa, Mythimna convecta, Nodaria cornicalis, Pandesma submurina, Plutella xylostella* and *Sceliodes cordalis* (Table 3). These species were chosen either because they are easily confused morphologically at the larval stages with *S. frugiperda*, often found as bycatch in regional pheromone traps or found colonising the same crops as *S. frugiperda*. Furthermore, 14 unidentified larvae from field sites were tested. Each sample was tested three times.

**Table 3.**
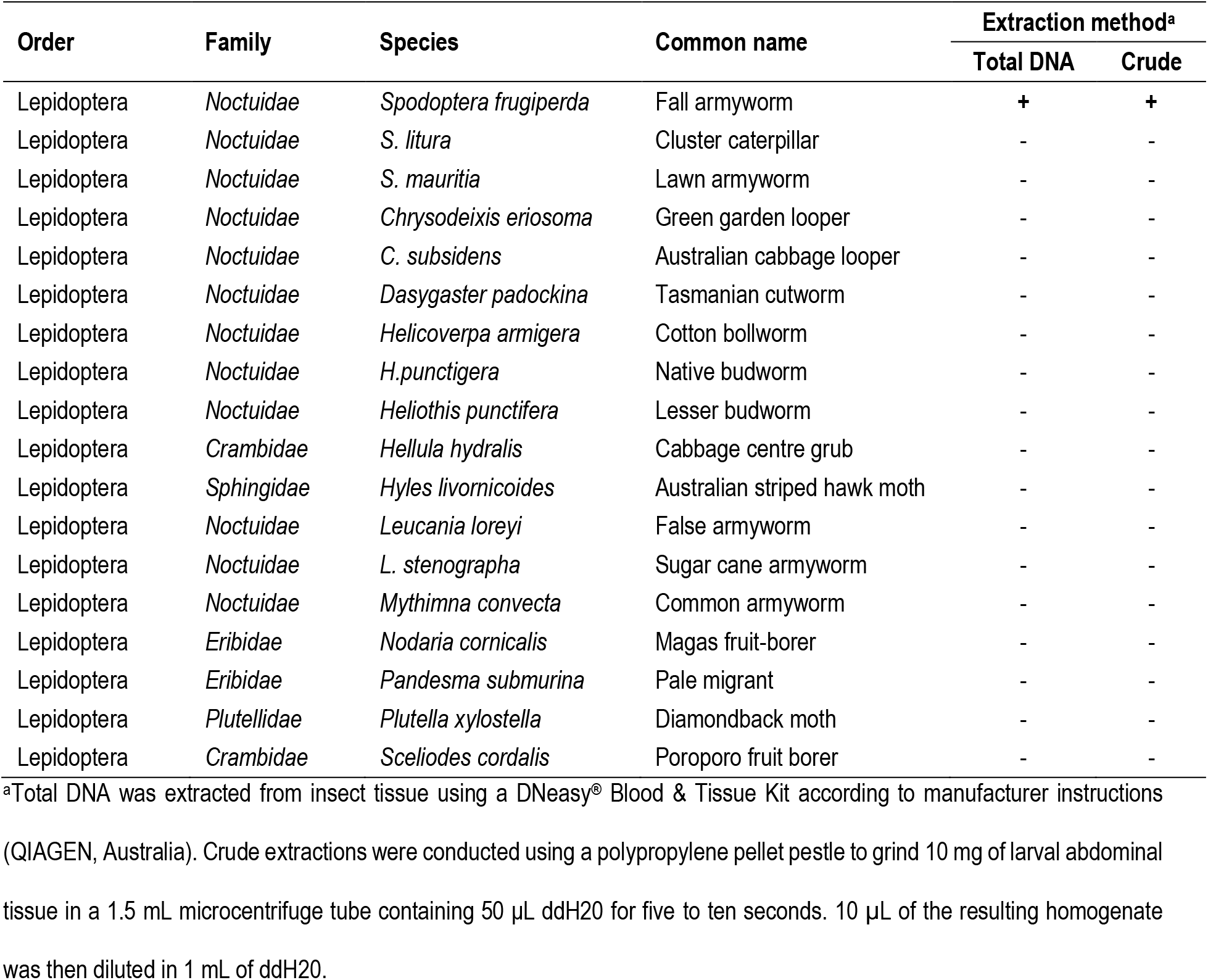
Specificity of *Spodoptera frugiperda* loop-mediated isothermal assay primer set FAW-1.2

To confirm the species of all larvae specimens tested, a region of the COX1 gene was amplified by PCR using GoTaq® G2 Green Master Mix (Promega, Australia) and primer set LepF (5’-TTCAACCAATCATAAAGATATTGG-3’) and LepR (5’-TAAACTTCTGGATGTCCAAAAAATCA-3’) or LCO1490 (5’-GGTCAACAAATCATAAAGATATTGG-3’) and HCO2198-puc (5’-TAAACTTCWGGRTGWCCAAARAATC-3’) (Coeur d’acier et al. 2014). The resulting amplicons were Sanger sequenced by the Australian Genome Research Facility (Perth, Australia).

*In silico* specificity analysis was performed against 12 congeneric species by aligning the nucleotide sequences of the COX1 region of each species with the *S. frugiperda* sequence amplified by primer set FAW-1.2.

## Results

### Parameter selection

For the *S. frugiperda* LAMP assay, the conditions that gave the fastest mean time to 10,000 flourescence (13.6 minutes) and most accurate detection (no type I or II errors) were 65°C amplification temperature with 18.5 pmol total primer concentration. These conditions also produced the second highest mean peak flourescence (48,157). Under these conditions, flourescence always reached 10,000 within 11 minutes for total DNA and 20 minutes for crude extractions. Annealing temperatures varied by 0.3°C at most. These parameters were used for the rest of the study.

### Diagnostic sensitivity and specificity

Testing of two known crude extraction *S. frugiperda* positives and two known crude extraction negatives (*H. punctigera* and *P. xylostella*) in duplicates 18 times each (72 reactions total each) resulted in no type I or type II errors. The average for time to 10,000 flourescence was 15.1 minutes (±0.23 SE), peak flourescence 53,000 (±956 SE) and annealing temperature 83.4°C (±0.04 SE, see Figure 1 for a typical reaction). Mean peak flourescence for the Genie II® was 49,784 (48 reactions) and for the Genie III®59,432 (24 reactions). The distribution of values for these metrics is shown in the boxplots in Figure 2.

**Figure 1.**
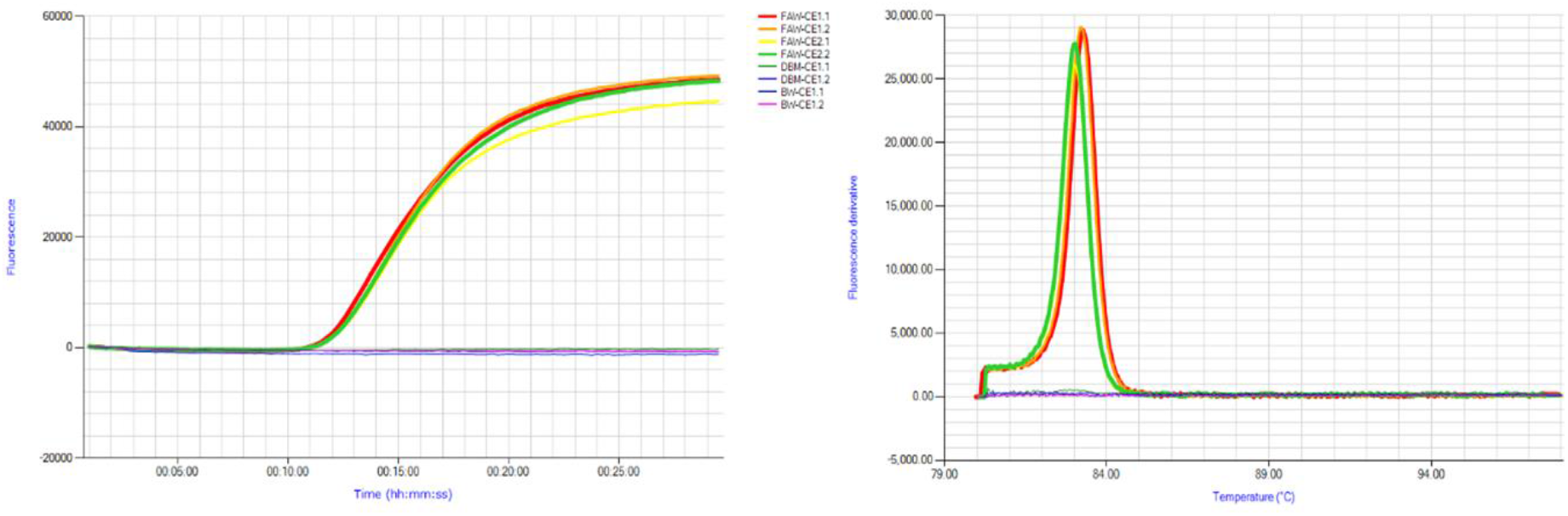
Typical amplification and annealing graphs obtained from Genie® instruments using primer set FAW-1.2 on crude extractions of *Spodoptera fruiperda* (FAW-CE), *Plutella xylostella* (DBM-CE) and *Helicoverpa punctigera* (BW-CE).

**Figure 2.**
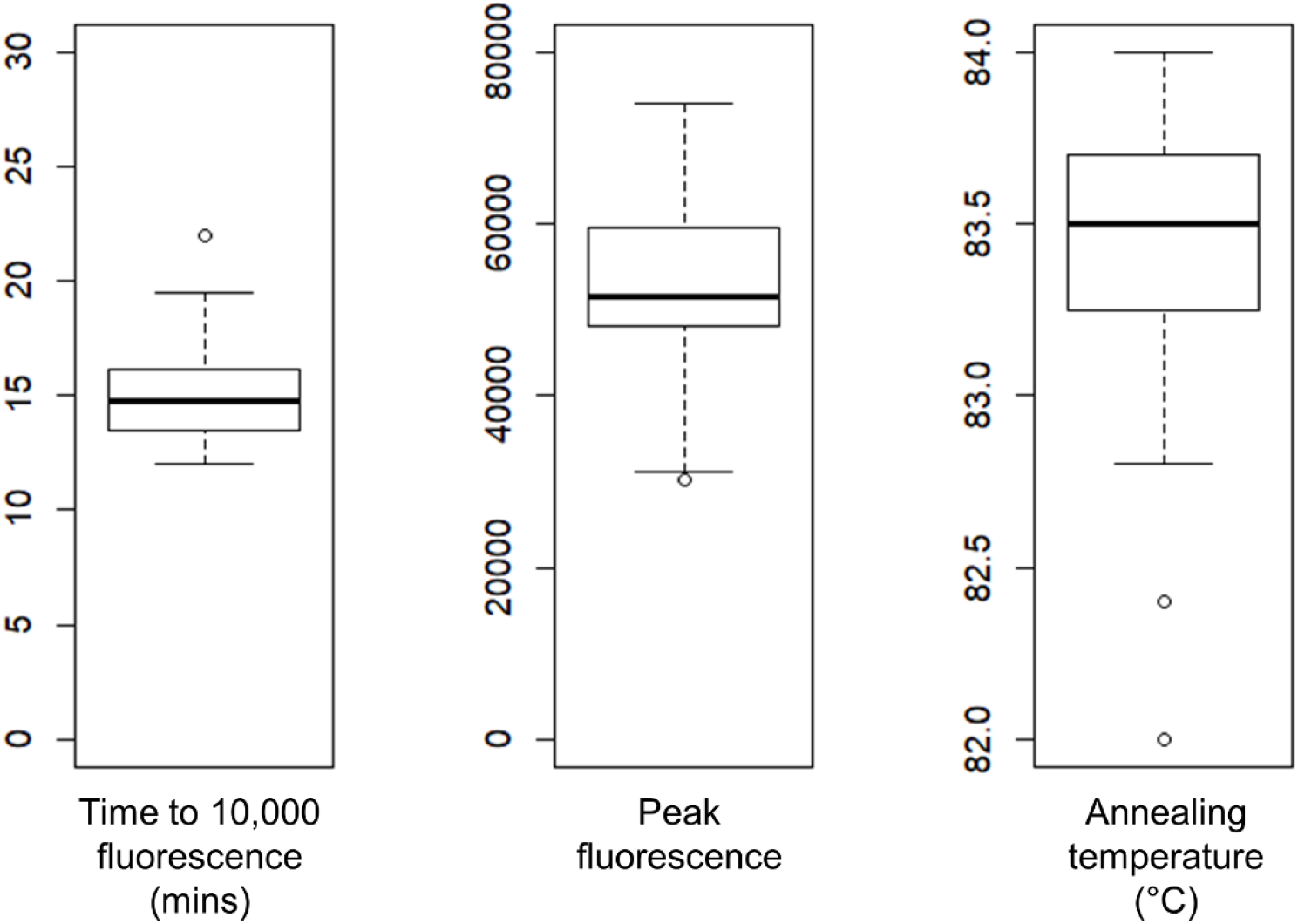
Boxplot statistics of 72 loop-mediated isothermal amplification (LAMP) reactions using the *Spodoptera frugiperda* primer set FAW-1.2 run on known positive crude extraction samples.

### Analytical sensitivity

The detection limit of the assay when testing three total DNA extractions of *S. frugiperda* with 37, 240 and 290 ng/μL was 37, 24 and 29 pg total DNA, respectively. Time to 10,000 fluorescence increased with decreasing DNA concentration with the 24 to 37 pg concentrations reaching 10,000 fluorescence between 19.5 and 30 minutes with equal or higher peak fluorescence after 50 minutes of amplification compared to dilutions with higher DNA concentrations. There was no amplification of 3.7, 2.4 and 2.9 pg concentrations after 1 h. Annealing temperatures varied no more than 0.2°C.

### Analytical specificity

Primer set FAW-1.2 did not cross react with total DNA or crude extractions of any of the other species tested (Table 3) and it correctly identified eight *S. frugiperda* larvae amongst 14 unidentified larvae. Furthermore, *in silico* analysis indicated that cross reaction with 12 congeneric species is unlikely with 4 to 10% differences from *S. frugiperda* in nucleotide sequences across the 194 bp primer region including the critical last three nucleotides of the primer binding regions (Figure 3).

**Figure 3.**
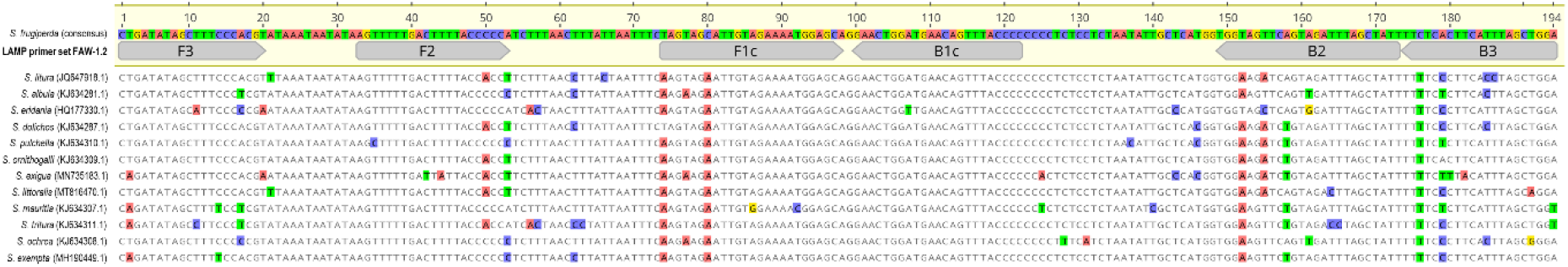
*In silico* analysis of *Spodoptera frugiperda* loop-mediated isothermal amplification (LAMP) primer set FAW-1.2 specificity against 12 congeneric species.

## Discussion

In this study, a sensitive, specific and in-field capable LAMP assay to detect *S. frugiperda* larvae was developed. A primer set based on a highly conserved region of the COX1 gene provided detection in total DNA and crude extraction within 30 min. The crude extraction technique is simple and rapid without requiring heat blocks, centrifuges or special buffers. The primer set did not cross-react with any other lepidopteran species tested and produced no type I or type II errors across 72 reactions of both known crude extraction positives and negatives. Furthermore, the primer set was able to detect 24 to 37 pg of DNA within 30 minutes indicating good sensitivity compared to similar assays (e.g. Przybylska et al. 2015). Therefore, this assay can be used in-field to quickly and accurately identify the presence of *S. frugiperda* at larval stages and thereby greatly assist with management decisions.

For field application purposes, the sensitivity and negligible false negative rate of the assay coupled with testing samples in duplicates will greatly diminish the likelihood of misdiagnosis of a larval population of *S. frugiperda* in crops. *Vice versa*, the high level of specificity and negligible false positive rate will prevent other species being diagnosed as *S. frugiperda*. These characteristics will ensure that correct management decisions will be made, especially concerning the selection and timing of insecticide applications. Combining the initial detection assay with in-field tests for insecticide resistance would further aid in selection of appropriate chemistries for *S. frugiperda* control. Further in-field validation of this assay in crops impacted by a diverse range of lepidopteran species would further improve confidence in using it as a decision support tool for growers. In a field laboratory context, provisions of standardised assay protocols and controls, and protocols for transport, storage and handling of consumables to prevent reagent degradation and sample contamination are also required. In cases where testing ambiguities or other technical errors occur, access to a centralised laboratory which can re-test diluted crude extractions or conduct total DNA extraction from larval samples is necessary.

The LAMP assay developed here is a viable option for rapid and affordable in-field molecular identification of *S. frugiperda* in Australia and may prove similarly useful in other world regions following validation against other regional-specific lepidopteran species.

## Acknowledgments

This work was funded by a DPIRD Building Grains project. All experiments were conducted using the DPIRD laboratory facilities in South Perth. We thank C. Brumley, A. Szito and A. Balfour-Cunningham for provision of larvae, and J. Baulch for technical support.

